# Role of non-coding RNA hsa_circ_0001495 in 16HBE cellular inflammation induced by PM_2.5_ and O_3_ combined exposure

**DOI:** 10.1101/2024.03.14.583416

**Authors:** HongJie Wang, Yi Tan, CaiXia Li, WenJia Jin, Ying Yu, Xuan Mu, XiaoWu Peng

## Abstract

**Background:** PM_2.5_ and O_3_ are the main air pollutants in China, and inflammation of the respiratory system is one of their main toxic effects. Cyclic RNAs are involved in many pathophysiological processes, but their relationship to the combined exposure to PM_2.5_ and O_3_ has not yet been investigated.

**Objective:** To elucidate the biological function played by hsa_circ_0001495 in the induction of 16HBE cellular inflammation by combined exposure to PM_2.5_ and O_3_.

**Method:** Detection of cell survival after 24h exposure of 16HBE cells to a combination of PM_2.5_ and O_3_ by CCK8. RT-qPCR and ELISA were used to detect inflammatory factors in 16HBE cells after co-exposing to PM_2.5_ and O_3_. CircRNA was screened using high throughput sequencing and bioinformatics analysis approaches. RNaseR experiments were carried out to verify the circular RNA properties of the circRNAs. Cytoplasmic-nuclear subcellular localisation assays and fish assays were used to verify the distribution of circRNAs in the nucleus versus the cytoplasm of the cell. To validate functions related with circRNA,RT-qPCR and ELISA were employed.

**Result:** Combined exposure to PM_2.5_ and O_3_ resulted in decreased cell viability.Combined exposure to PM_2.5_ and O_3_ resulted in 16HBE inflammation. High throughput sequencing and RT-qPCR results showed that the expression of hsa_circ_0001495 was significantly downregulated in 16HBE exposed to PM_2.5_ and O_3_ in combination. Hsa_circ_0001495 is not easily digested by RNaseR enzymes and has the properties of a circular RNA. Hsa_circ_0001495 is expressed in the cytoplasm as well as in the nucleus, but its distribution is predominantly in the cytoplasm.

**Conclusion:** In 16HBE cells, combined exposure to PM_2.5_ and O_3_ can induce an inflammatory response.hsa_circ_0001495 plays an inhibitory role in the inflammatory response of 16HBE cells that can be induced by combined exposure to PM_2.5_ and O_3_.

## 1. Introduction

PM_2.5_ is mainly generated through natural pathways such as volcanic eruptions and forest fires ^[1]^and anthropogenic pathways such as fossil fuel combustion due to human production and living ^[2]^ Numerous epidemiological studies have shown ^[3]^ that living in an environment with high concentrations of PM_2.5_ for a long period of time can significantly increase the incidence of disease and mortality in the population, such as respiratory and cardiovascular diseases and many other diseases. It has been reported ^[4]^ that for every 10 ppb (21.44 μg/m^3^) increase in O_3_ concentration, the respiratory mortality rate increases by 0.64% (95% PI: 0.31%-0.98%). The association between atmospheric O_3_ pollution and increased risk of respiratory disease is well established^[5, 6]^, and inhalation of O_3_ may damage lung epithelial cells ^[7, 8]^.Wong ^[9]^ et al. found that the combined exposure to ultrafine particles and O_3_ increased the extent of lung damage, with severe damage to both the large airways and the small airways, and was not a additive effect of a single pollutant exposure PM_2.5_ and O_3_ and the inflammatory effect on the respiratory system is one of the main toxic effects, and there is a synergistic effect between the two. A large number of studies have shown that PM_2.5_ and O_3_ and the mechanism of respiratory system damage caused by the inflammatory response is considered to be the basic pathogenic mechanism. Happo ^[10]^ and other studies have found that PM_2.5_ can directly stimulate alveolar macrophages (AMs) to secrete a large number of pro-inflammatory cytokines, chemokines, leading to diffuse inflammation in the lungs. It was found ^[11]^hat IL-1α released from O_3_-induced tissue damage and inflammation is mediated by MyD88 signalling in epithelial cells. In recent years, with the rapid development of biological science theories and technologies, circular RNA (circRNA) has received keen attention from research scholars. circRNA is a closed-loop non-coding RNA covalently linked at the 3’ and 5’ ends produced in the process of reverse splicing, and it has a high degree of tissue-expression specificity in the eukaryotic transcriptome^[12]^. In this experimental study, we found that an abnormally low expression of has_circ_0007766 occurred after compound exposure of human bronchial epithelial cells (16HBE) stained with PM_2.5_ (100 μg/ml) + O_3_ (300ppb,2h).This study was designed to explore the inflammatory response induced by compound exposure of 16HBE cells with PM_2.5_ and O_3_ and the has_circ_0007766 in this process to provide a scientific basis for PM_2.5_-induced respiratory diseases.

## 2. Materials and methods

### 2.1 Cell and cell culture

Human bronchial epithelial cells (16HBE) were obtained from the group of Wu Jianjun from Guangzhou Medical University, China. 16HBE cells were grown in MEM complete medium (Cienry,China), which consisted of 1% penicillin/streptomycin (gibico, USA), 5% foetal bovine serum (Four Seasons Green, China), and 94% MEM basal medium, and cultured at air-liquid interface with Transwell (Corning, USA) at a temperature of 37°C and CO_2_ concentration of 5%. The cells were cultured at the air-liquid interface using Transwell (Corning, USA) at a temperature of 37°C and a concentration of 5% carbon dioxide.

### 2.2 Preparation of PM_2.5_ and O_3_ subjects

Form March to November 2021,On the roof of an office building (Huangpu District, Guangzhou, Guangdong) collecting PM_2.5._ The TH1000C large flow sampler (Wuhan Tianhong, China) was used to collect PM2.5 with a sample flow of 1.05 m^3^/min and 72h of continuous sampling. O_3_ occurs in ozone calibrators(2Btech,USA).

### 2.3 Cellular contamination

The Cloud System Particulate Dyeing Instrument(VOTROCELL CLOUD12, Germany) as well as the Continuous Fluid External Exposure System(VOTROCELL 12/12, Germany) were used to compound exposure to PM_2.5_ and O_3_ in 16HBE cells.

### 2.4 Cell Counting Kit-8 Assay

The Cell Counting Kit-8 (CCK-8) (Biosharp, China) was used to measure cell viability. Human bronchial epithelial cells were cultured in a 12-well Transwell and treated with PM_2.5_ and O_3_ for 24 hours, and the grouping of contamination was divided into 0, PM_2.5_ (100 μg/ml), PM_2.5_ (100 μg/ml) + O_3_ (300ppb,2h), PM_2.5_ concentrations were selected based on previous experiments conducted by the group and relevant literature. ^[13-15]^, and the concentration of O_3_ toxicity was taken from the literature ^[16-18]^,and the experiment was carried out in three biological replicates. After 24h of contamination, the cells were washed with PBS, added with CCK8 reagent mixture, and incubated for 1h, protected from light. Absorbance at 450nm was measured by microplate reader(BioTek Synergy HT,USA). Cell viability was calculated according to the instructions.

### 2.5 RT-qPCR analysis

The experimental groupings were: 0, PM2.5 (100 μg/ml), PM_2.5_ (100 μg/ml) + O_3_ (300ppb,2h). Human bronchial epithelial cells were inoculated into 12-well Transwell cell chambers at 1.5×105 cells/well and cultured for 24h after exposure to a combination of PM_2.5_ and O_3_. SevenFast® Total RNA Extraction Kit(SEVEV,China) was used to extract total cellular RNA. Evo M-MLV RT Kit with gDNA Clean for qPCR II (Accurate Biology, China) was used to reverse transcribe the total RNA. qRT-PCR was conducted using SYBR®Green Pro Tap HS(Accurate Biology,China)and detected by Step One Plus(Applied Biosystems,USA). Relative gene expression level = 2-ΔΔCt.ACTB (β-actin) was used as an internal reference gene for correction of relative gene expression. The primers are shown in Table S1.

### 2.5 Enzyme-linked immunosorbent assay

The experimental groupings were: 0, PM_2.5_ (100 μg/ml), PM_2.5_ (100 μg/ml) + O_3_ (300ppb,2h). Human bronchial epithelial cells were inoculated into 12-well Transwell cell chambers at 1.5×10 ^5^ cells/well and cultured for 24h after exposure to a combination of PM_2.5_ and O_3_. ELISA kits were used to detect the levels of cellular inflammatory factors IL-8, IL-1β protein exp(Jiubang,China)ression. The OD value of the measured standard was taken as the horizontal coordinate, and the concentration value of the standard (the concentration of the standard was 80, 40, 20, 10, 5, 2.5 pg/mL in order) was taken as the vertical coordinate, and the standard curve was plotted by Excel software, and a linear regression equation was obtained, and the OD value of the samples was substituted into the equation, and the concentration of the samples was calculated.

### 2.5 High-throughput sequencing and data analysis

Samples of 16HBE were collected for high-throughput sequencing after exposure to PM_2.5_+O_3_ (0, PM_2.5_+O_3_) for 24 hours.The technology has been done at Shanghai Kangcheng Biotechnology Co., Ltd.

### 2.6 RNase R experiment

Cells were inoculated into 12-well Transwell cell chambers at 1.5×10 ^5^ cells/well and placed in the incubator for 24 h. Total cell RNA was extracted after the cell fusion rate reached 70%-80%.Then the extracted cell RNA was subjected to the experiment according to the instructions of the de-novo enzyme experiment. Experimental grouping: RNase R+ group (RNase R enzyme treatment) and RNase R-group (no RNase R enzyme treatment), 4U of RNase R reagent was added to 2.4 μg of total RNA (RNase R+ group), and after incubation at 37°C for 10 min, reverse transcription as well as qRT-PCR experiments were performed to detect hsa_circ_ 0007766 and the expression of the internal reference ACTB (2-ΔΔCt). The internal reference ACTB in the RNase R-group was used for subsequent calculations. RNase R experiment were carried out according to the RNase R instructions (Guangzhou Geneseed Biological Technology Co., Ltd, China).

### 2.7 Nucleocytoplasmic separation experiment

Cells were inoculated with 10^6^ cells/well in a 10 cm dish and placed in an incubator for 24 h. After the cell fusion rate reached 70%-80%, nucleoplasmic separation experiments were carried out using the Invitrogen PARIS™ am1921 kit (Invitrogen AM1921, USA), and the nuclei and cytoplasm were extracted. qRT-PCR experiments were carried out to detect the expression (2^-Δ Δ Ct^) of the hsa_circ_0007766 and the internal reference genes (ACTB, U6), and the primer sequences are shown in Table S2.

### 2.8 Fluorescence in situ hybridization

The FISH procedure followed the RNA FISH kit instructions(GenePharma, China). Finally, cells were examined with IX71 Inverted Research System Microscope (Olympus, Japan). The FISH probe were designed by the Suzhou GenePharma Co., Ltd. The probe sequences are shown in Table S2.

### 2.9 Transient transfection of cyclic RNA

#### 2.9.1 Transfection efficiency assay

The 16HBE cells were inoculated in 6-well plates at a density of 4.5×10 ^5^ cells/well. 80%-90% cell density was achieved after 24h, and the next experiment could be carried out. The experimental groups were as follows: hsa_circ_0007766 group (hsa_circ_0007766 plasmid), negative transfection control group (NC), MOCK group (transfection reagent control group). Transfection complexes were prepared: part A: 121 μl Opti-MEM medium + 4 μl lipofectamine 3000; part B: 112.23 μl Opti-MEM medium + 6.77 μl overexpression solution + 5 μl P3000, part C: 114 μl Opti-MEM medium + 6 μl NC solution + 5 μl P3000; Part D: 120 μl Opti-MEM medium + 5 μl P3000. The transfection complex was prepared by mixing A and B, A and C, A and D in the ratio of 1:1, and placed at room temperature for 20 min, and then the transfection complexes were added into cell culture plates according to the experimental grouping, and the amount of solution per well was 250 μl of transfection complex + 1.75 ml Opti-MEM culture medium. After 5h of culture, the old medium was aspirated, the cells were rinsed using PBS buffer to remove the residual liquid, and then 2ml of MEM complete medium was added to culture the culture for 24h, then the cellular RNA was extracted, and the changes of hsa_circ_0007766 level were detected by qRT-PCR. The overexpression sequence was designed and synthesised by Suzhou Gemma.

#### 2.9.2 Validation of cyclic RNA function

Experimental grouping: 16HBE cells were divided into four groups, NC group (overexpression of plasmid-negative transfection sequence), hsa_circ_0007766 group (overexpression of plasmid by hsa_circ_0007766), NC+ PM_2.5_ (100 μg/ml) + O_3_ (300ppb, 2h) group (overexpression of plasmid-negative transfection sequence +PM_2.5_ (100 μg/ml) + O_3_ (300ppb, 2h)), hsa_circ_0007766+ PM_2.5_ (100 μg/ml) + O_3_ (300ppb, 2h) group (hsa_circ_0007766 overexpression plasmid +PM_2.5_(100μg/ml)+O_3_(300ppb, 2h)), the transfection complex was prepared as in 2.9.1, according to the experimental grouping, the transfection complex was added to the upper chamber of the Transwell with the Opti-MEM medium, and the lower chamber was added with 1.5 ml of the Opti-MEM medium, and then the waste liquid was discarded and washed with PBS for two times, and then replaced with the serum free medium of the MEM. Then PM2.5+O3 staining, after 24h of culture, the cell RNA was extracted, and then by qRT-PCR (reaction system and conditions are the same as 1.5.1), the gene expression=2-ΔΔCt; ELISA detected the changes in the transcription and protein levels of the inflammatory factors IL-1β and IL-8 (the results of the experiment were based on the OD value of the measured standard as the horizontal coordinate, the standard concentration value (the concentration of the standard in the order of: 80, 40, 20, 10, 5, 2.5 pg/mL) as the vertical coordinate, the standard curve was drawn by Excel software, and the linear regression equation was obtained, the OD value of the sample was substituted into the equation, and the concentration of the sample was calculated).

### 2.10 Statistical analysis

For statistical data analysis, GraphPad Prism 8.0 and SPSS 25.0 software were utilized. All studies were carried out three times, and the findings were reported as t mean standard deviation 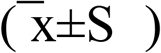. When comparing two groups, the T test was employed, and when comparing multiple groups, the one-way ANOVA was utilized. The difference was statistically significant when P<0.05 was used.

## 3 Result

### 3.1 16HBE cell activity after 24h exposure to PM2.5 and PM2.5+O3 complexes

After 24h of contamination,the cellular activity of the PM2.5 group was reduced (P<0.05) compared with the control group, and the reduction of cellular activity was more obvious in the PM2.5+O3 group (P<0.05); the cellular activity of the PM2.5+O3 concentration group was decreased (P<0.05) compared with that of the PM2.5 group(Figure 3.1).

**Figure 3.1.**
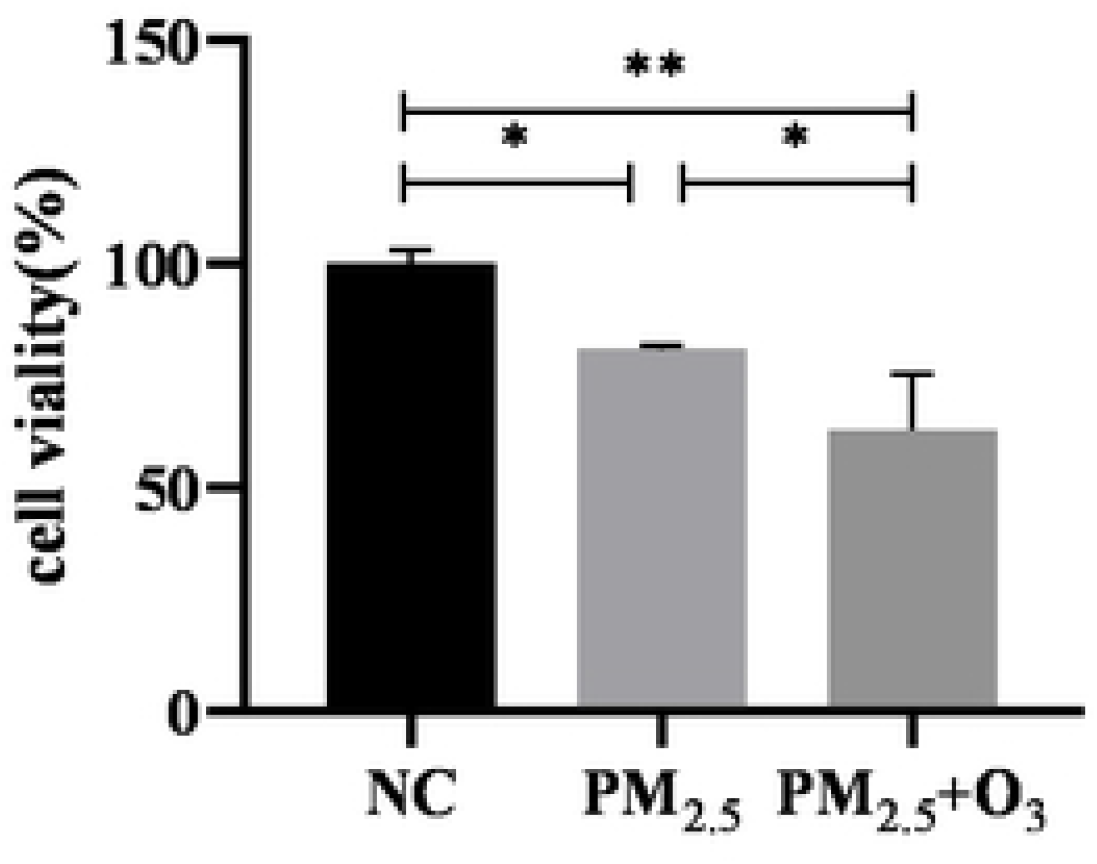
Effect of PM2.5 and PM2.5+O3 staining on the viability of 16HBE cells after 24h of poisoning

### 3.2 Inflammatory effects of 16HBE after 24h exposure to PM2.5 and PM2.5+O3 composite exposure

After 24 h of contamination, the expression of inflammatory factors IL-1β and IL-8 transcript and protein levels were elevated in the PM2.5 group compared with the control group (P < 0.05); the expression of inflammatory factors transcript and protein levels were more significantly elevated in the PM2.5+O3 group (P < 0.05); the expression of inflammatory factors IL-1β and IL-8 transcript and protein levels were elevated in the PM2.5+O3 concentration group compared with the PM2.5 group (P < 0.05). transcript and protein levels were elevated (P < 0.05) (Figure 3.2).

**Figure 3.2.**
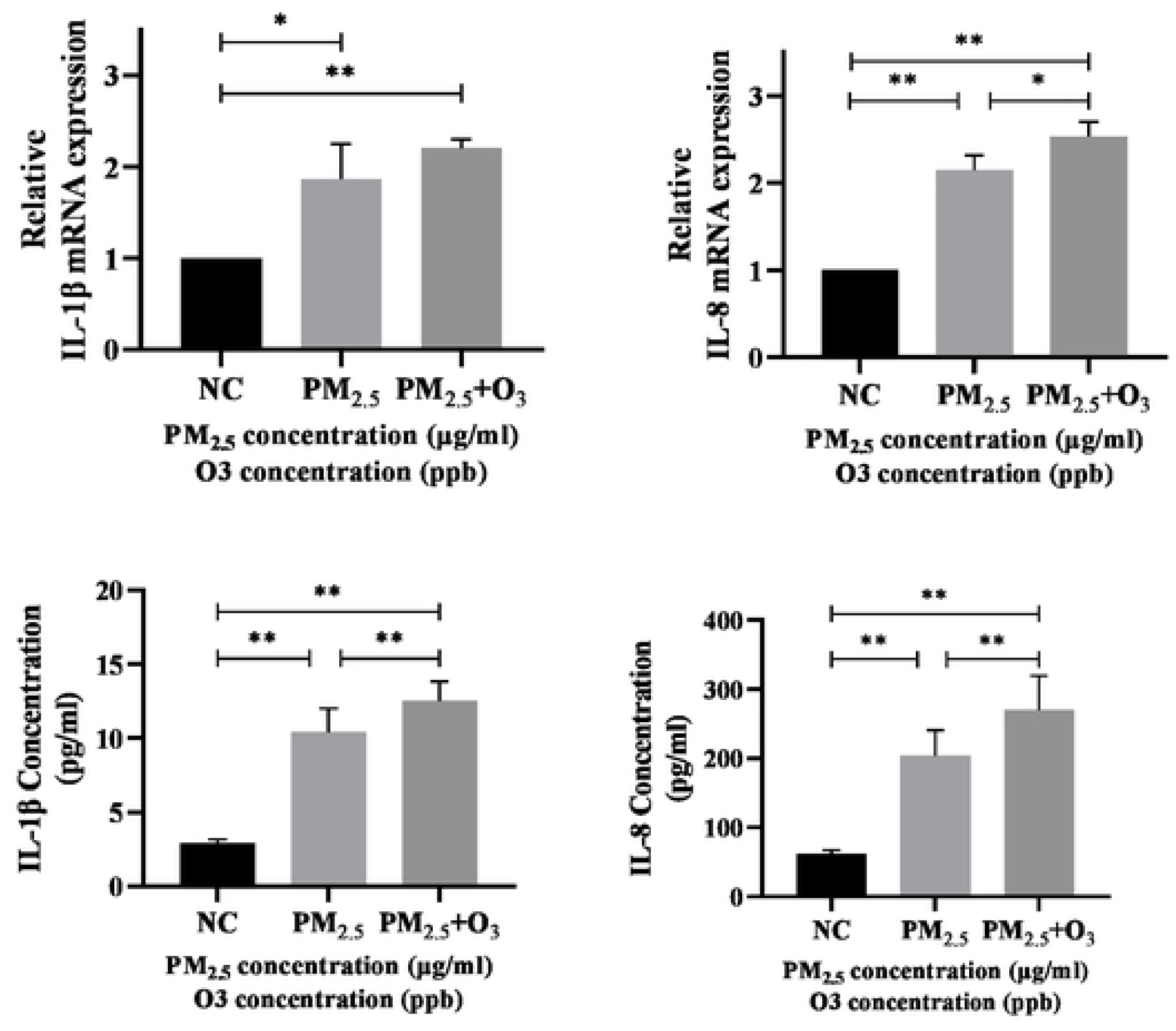
Transcription and protein expression levels of cellular inflammatory factors IL-1β and IL-8 after 24 h of PM2.5 and PM2.5+O3 stimulation

### 3.3 qRT-PCR validation of circRNAs associated with cellular inflammatory effects

Based on the high-throughput sequencing results, 10 differentially expressed circRNAs associated with cellular inflammatory pathways were screened out, and in order to verify the accuracy of the RNA-seq results, we performed qRT-PCR on these 10 circRNAs again. Hsa_circ_0001495 expression level was consistent with the sequencing results, and hsa_circ_0001495 was identified as the target circRNA. After staining 16HBE cells with 0, PM2.5 group, and PM2.5+O3 composite exposure group, respectively, for 24 h, the hsa_circ_0007766 expression level was reduced in PM2.5 group as well as in PM2.5+O3 composite exposure group compared with the control group (P < 0.05), and the PM2.5+O3 composite exposure group had a more pronounced (P < 0.05) reduction in hsa_circ_ 0001495ession level was reduced in the PM2.5+O3 compound exposure group compared with the PM2.5 group (P < 0.05). It indicates that hsa_circ_0001495 suppresses the inflammatory response in 16HBE cells.(Figure 3.3).

**Figure 3.3.**
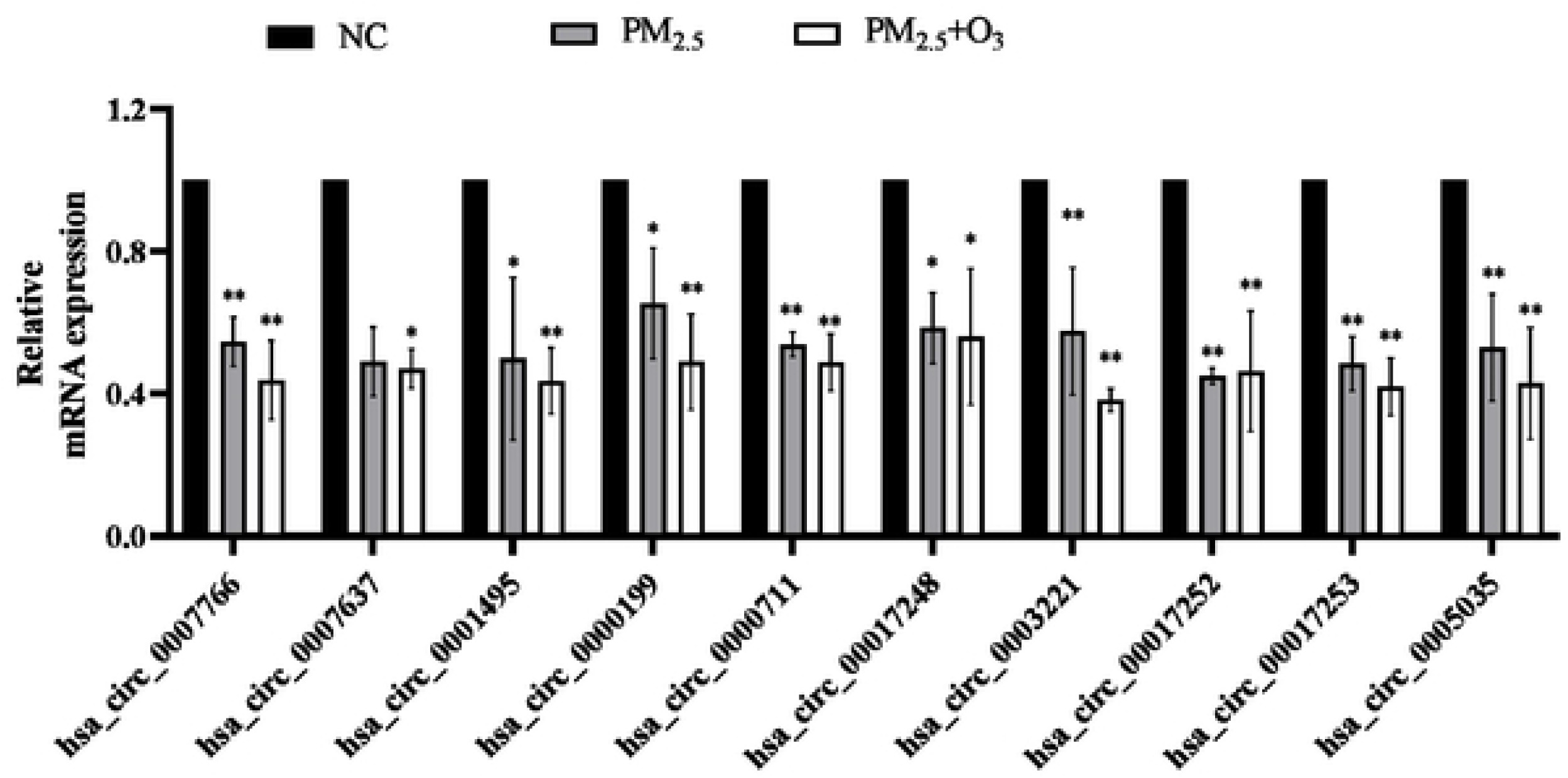
mRNA transcript levels of circRNAs in PM2.5 and PM2.5+O3-stained 16HBE cells after 24h

### 3.4 RNase R digestive test

After RNase R treatment, the expression of the endogenous ACTB was significantly reduced (P < 0.05), whereas hsa_circ_0001495 was able to tolerate RNase R digestion, suggesting that hsa_circ_0001495 has a cyclic structure. (figure 3.4)

**Figure 3.4.**
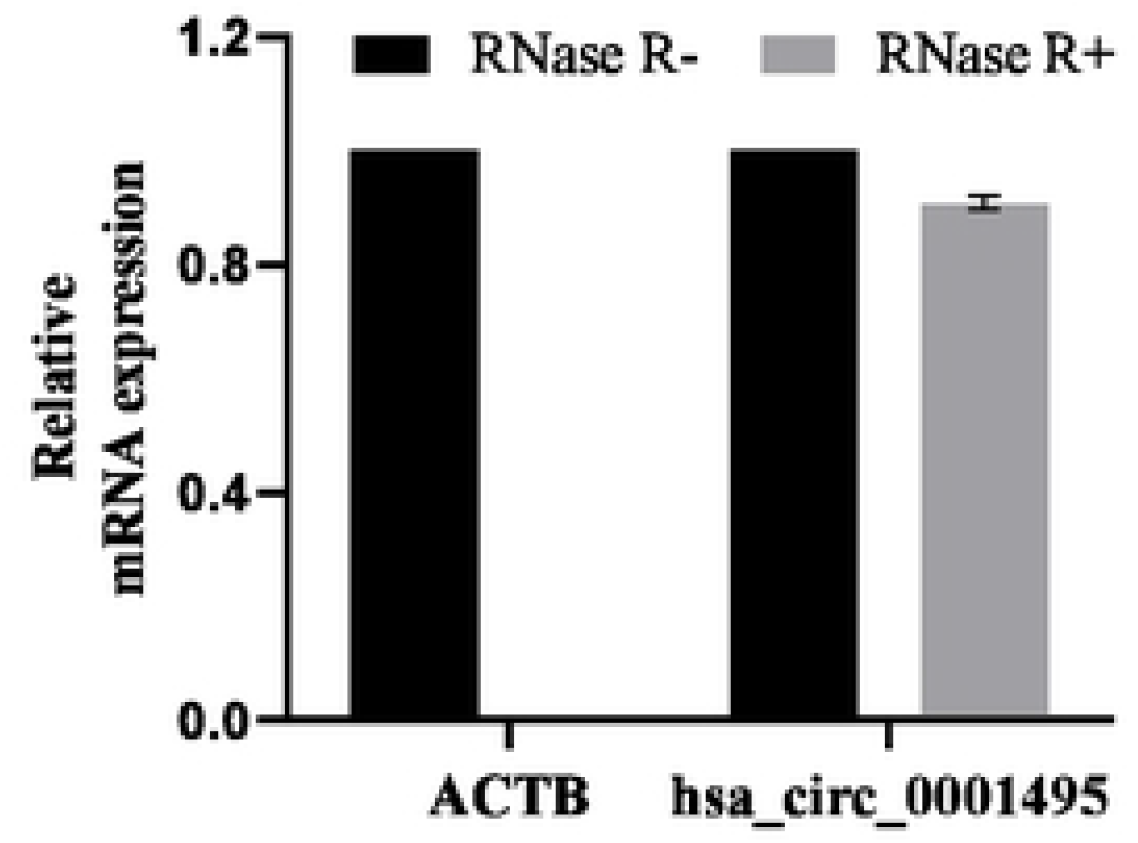
RNaseR digestion experiment

### 3.5 Nucleocytoplasmic separation experiment

ACTB and U6 were mainly distributed in the cytoplasm and nucleus, respectively, indicating that the nucleoplasmic segregation experiment was successful, while hsa_circ_0001495 was mainly expressed in the cytoplasm. (figure 3.5)

**Figure 3.5.**
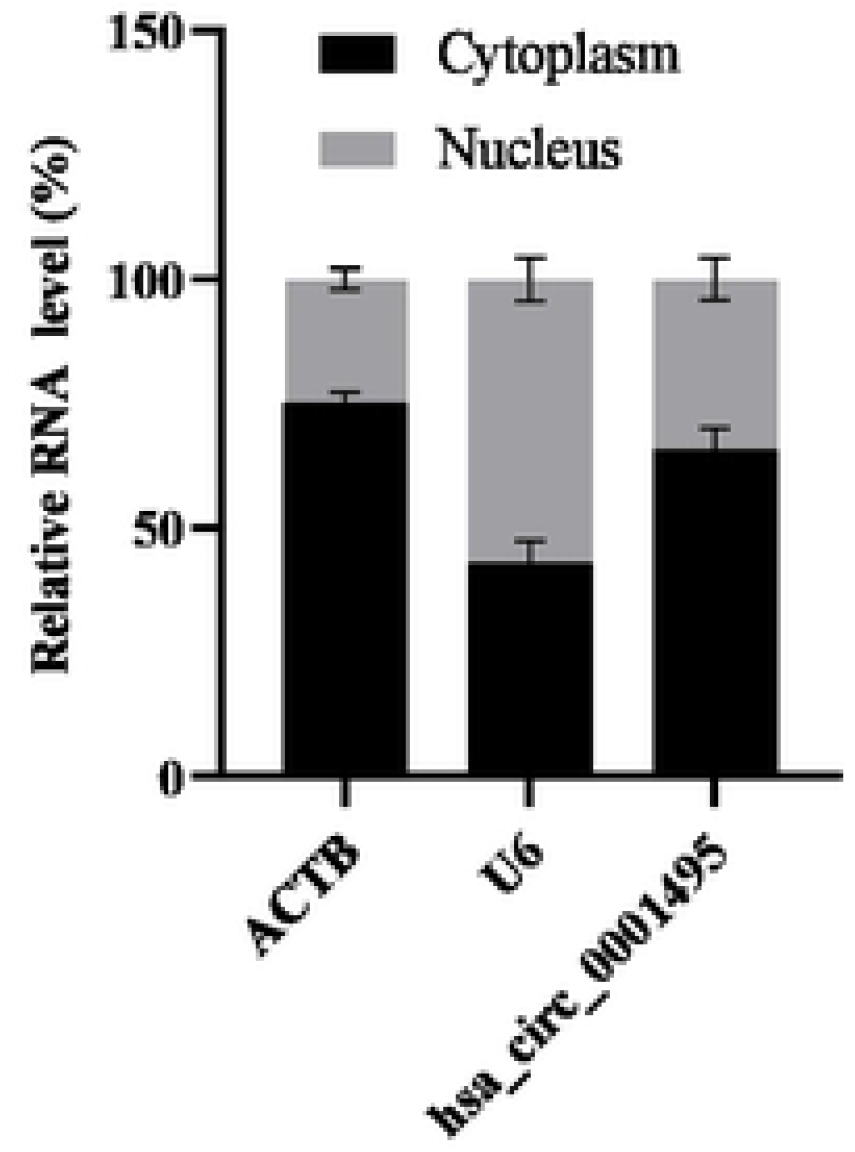
Nucleocytoplasmic separation experiment

### 3.6 Fluorescence in situ hybridization experiment

To further verify the distribution of hsa_circ_0001495 in cells, we localized hsa_circ_0001495 through fluorescence in situ hybridization experiment, and found that Cy3-labeled hsa_circ_0001495 was distributed in both nucleus and cytoplasm (Figure 3.6)

**Figure 3.6.**
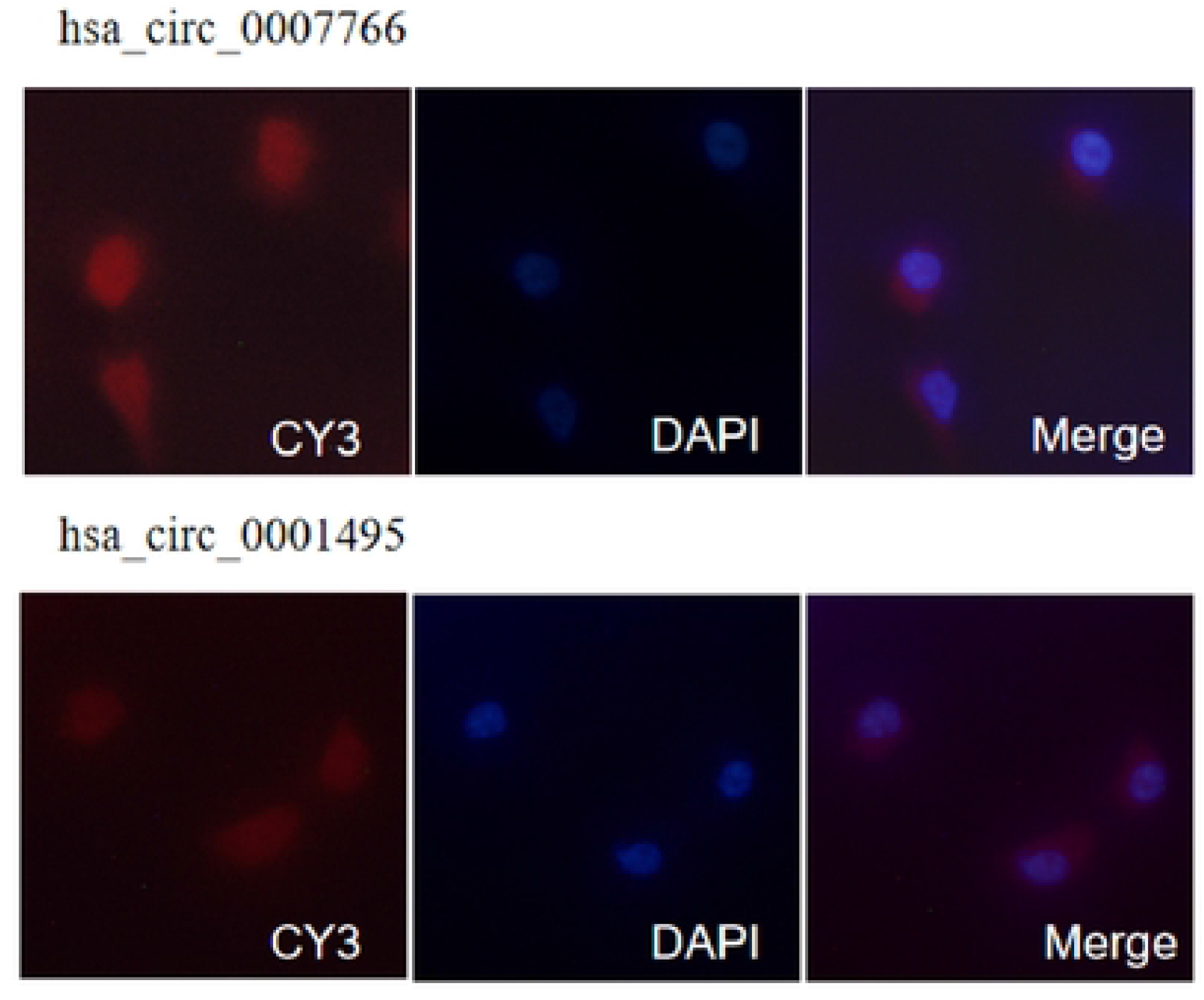
Fluorescence in situ hybridization experiment

### 3.7 hsa_circ_0001495 Overexpression effect

Compared with the NC group, the hsa_circ_0007766 overexpression group was able to effectively increase the expression of hsa_circ_0007766 (P < 0.05). (Figure 3.7)

**Figure 3.7.**
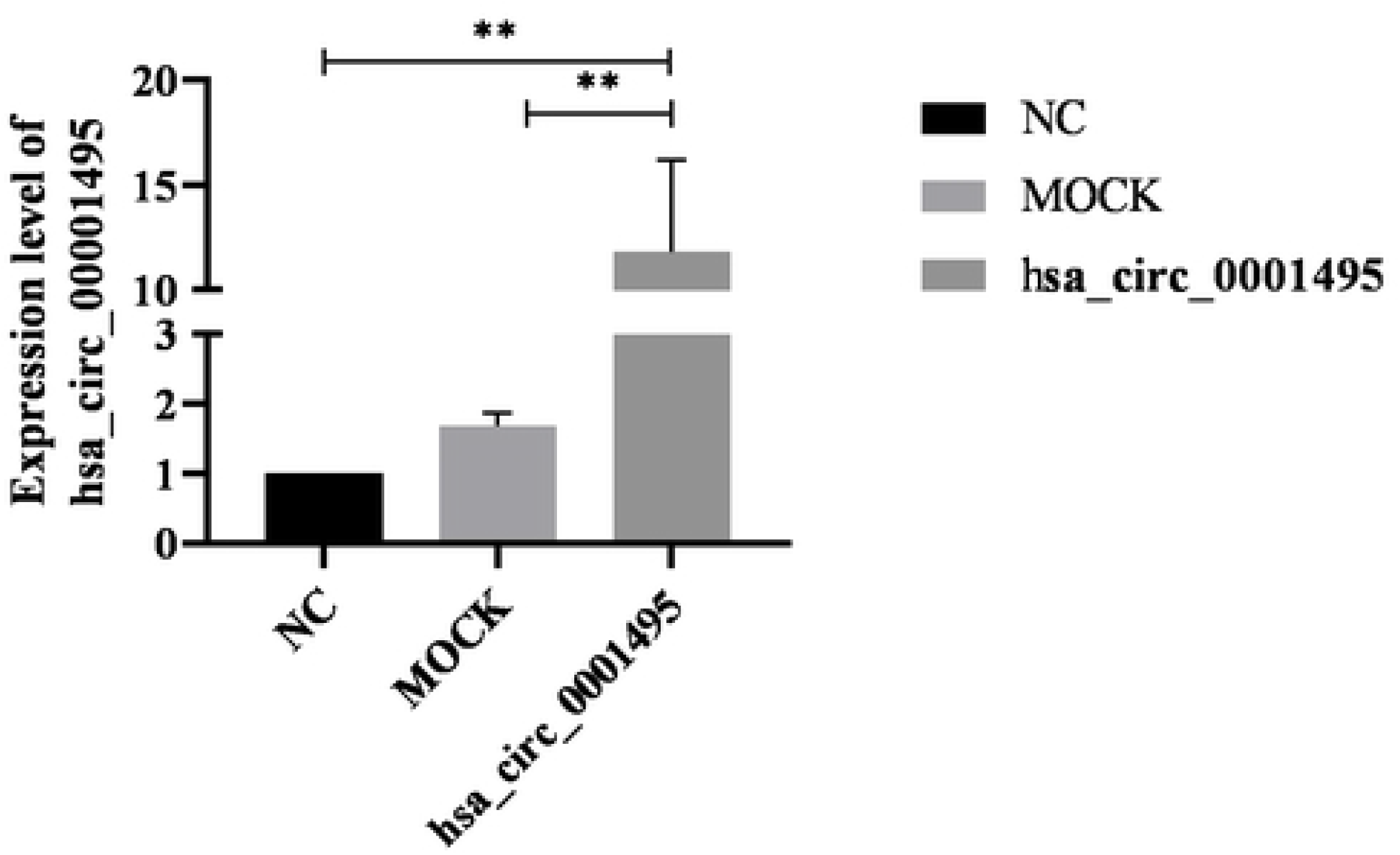
Hsa_circ_0001495 Overexpression effect

### 3.8 Hsa_circ_0001495 functional verification experiment

The cellular inflammatory effect was elevated after PM2.5+O3 complex exposure to 16HBE cells (P < 0.05); compared to the NC group, the cellular inflammatory effect was decreased after overexpression of hsa_circ_0007766 (P < 0.05); compared to the NC+PM2.5 group, the cellular inflammation in the hsa_circ_0007766+PM2.5 group effect was decreased (P < 0.05). Thus, overexpression of hsa_circ_0007766 decreased the cellular inflammatory effect, suggesting that hsa_circ_0007766 has an anti-inflammatory effect in 16HBE cell inflammation caused by PM2.5+O3 compound exposure. (Figure 3.8)

**Figure 3.8.**
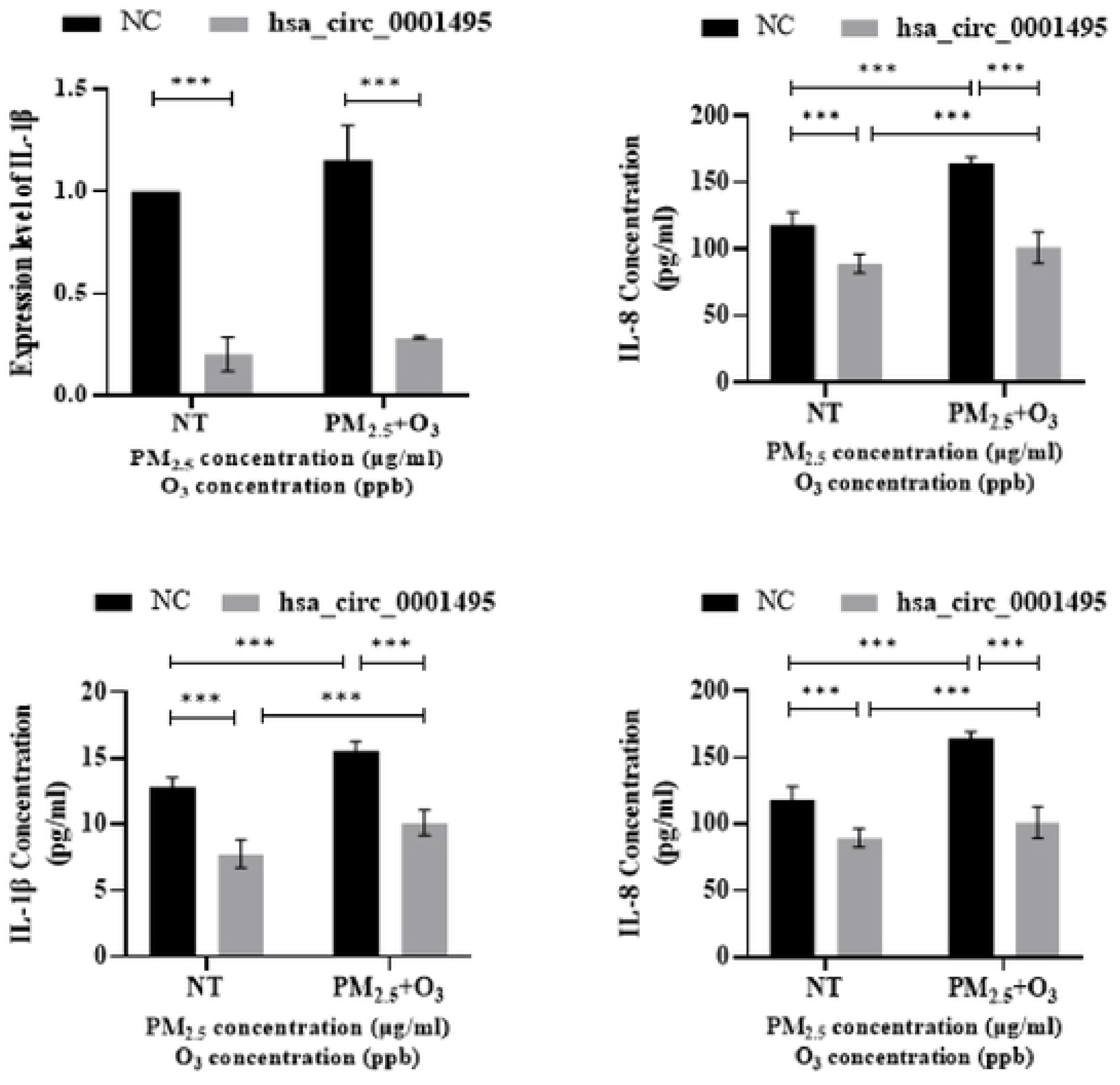
Transcription and protein expression levels of IL-1β, IL-8 after overexpression of hsa_circ_0007766 or combined exposure to PM2.5+O3

## 4 Discussion

According to the 2022 China Ecological Environment Status Bulletin, the average atmospheric PM2.5 and O3 concentrations in 339 cities at prefecture level and above in China were 29 μg/m^3^ and 145 μg/m^3^, respectively, and the number of exceedance days in which PM2.5 and O3 were the primary pollutants accounted for 36.9% and 47.9% of the total number of exceedance days, respectively; compared with 2021, the proportion of exceedance days of both PM2.5 and O3 increased, and the concentration of O3 increased. Compared with 2021, the proportion of days with exceedance of atmospheric PM2.5 and O3 increased, and the concentration of O3 increased. Short-term exposure to PM2.5 and warm-season O3 was significantly associated with an increased risk of mortality^[19]^,and the combined PM2.5 and O3 pollution has become a major air pollution problem in China. Many epidemiological evidences show that PM2.5 is closely related to respiratory diseases, and the damage of fine particulate matter to the respiratory system has been confirmed in vivo and in vitro. The respiratory system is the main route of PM2.5 inhalation^[20]^ Current epidemiological data show a significant correlation between PM2.5 and respiratory diseases. Long-term exposure to fine particulate matter (PM2.5) is associated with reduced lung function in adults^[5, 21]^.

The association between atmospheric O3 pollution and increased risk of respiratory diseases is well established^[6]^. Inhalation of O3 may damage lung epithelial cells^[7, 8]^.The results of Kim ^[22]^ and others have shown that long-term exposure to O3 is associated with an increased risk of respiratory mortality. Long-term standards for O3 and PM should be considered to protect the respiratory health of the general population and patients with chronic respiratory diseases.

Inflammation is a defence-based pathological response following an external stimulus, and excessive inflammation is considered to be the main causative event leading to the development or exacerbation of respiratory diseases ^[23]^.The inflammatory effect of PM2.5 and O3 on the respiratory system is one of their main toxic effects, and there is a synergistic effect between them. Numerous studies have shown that the inflammatory response is considered to be the underlying pathogenic mechanism in the mechanism of respiratory system damage caused by PM_2.5_ and O_3_. Sokolowska^[24]^ and others reviewed the latest data on the mechanisms of O_3_ damage to different cell types and pathways, with a focus on the role of the IL-1 family of cytokines and the related IL-33. It has been suggested ^[25]^ that the IL-33/ST2 pathway contributes to O_3_-induced airway hyperresponsiveness in male mice, and that the interaction of O_3_ and traffic-associated PM_2.5_ produces significantly more hydroxyl radicals than PM_2.5_ alone, suggesting that combined exposure to PM_2.5_ and O_3_ is more likely to lead to organismal inflammation. Therefore, it is necessary to carry out compound exposure studies to provide an important scientific basis for the prevention and control of environmentally related diseases.

So far, although O_3_ and PM_2.5_-induced respiratory inflammation has received much attention from many scholars, the molecular mechanism of compound exposure is still unclear, and there is an urgent need to find new ways to explore. A large number of circRNAs have been found to regulate gene expression and play important biological functions. circRNAs play an important role in body inflammation. Therefore, we explored the role of circRNAs in the inflammatory effects caused by O_3_ and PM_2.5_ exposure in vitro, aiming to provide gene therapy targets for O_3_ and PM2.5-induced inflammation at the level of non-coding RNAs, and to expand a new direction for the study of inflammatory mechanisms.

In this study, we identified for the first time the relationship between hsa_circ_0007766 and the inflammatory response of 16HBE cells induced by the combined exposure of PM_2.5_ and O_3_. hsa_circ_0007766 was abnormally low expressed in the inflammatory cells, revealing its biological function as an inhibitor of inflammation in the inflammatory response of 16HBE cells induced by the combined exposure of PM_2.5_ and O_3_. These findings provide new ideas and directions for the search of diagnostic and therapeutic targets for inflammation induced by the combined exposure of PM_2.5_ and O_3_.

## Acknowledgements

This work was supported by the National Natural Science Foundation of China(Grant no.21477045),the National Key Research and Development Program of China (No.2023YFC39005204),the Central Public-Interest Scientific Institution Basal Research Fund(Grant no.PM-zx703-202004-155,Grant no.PM-zx703-202204-164).

## Tables

**TableS1.**
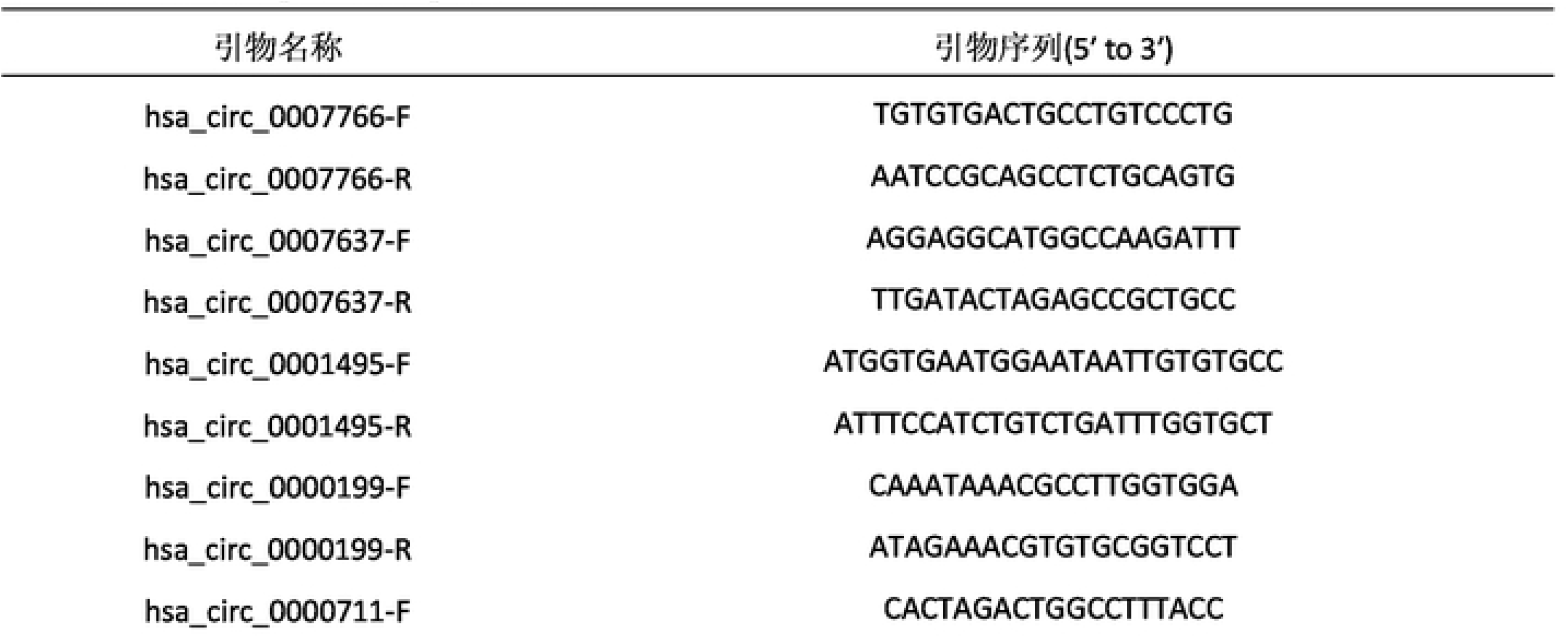
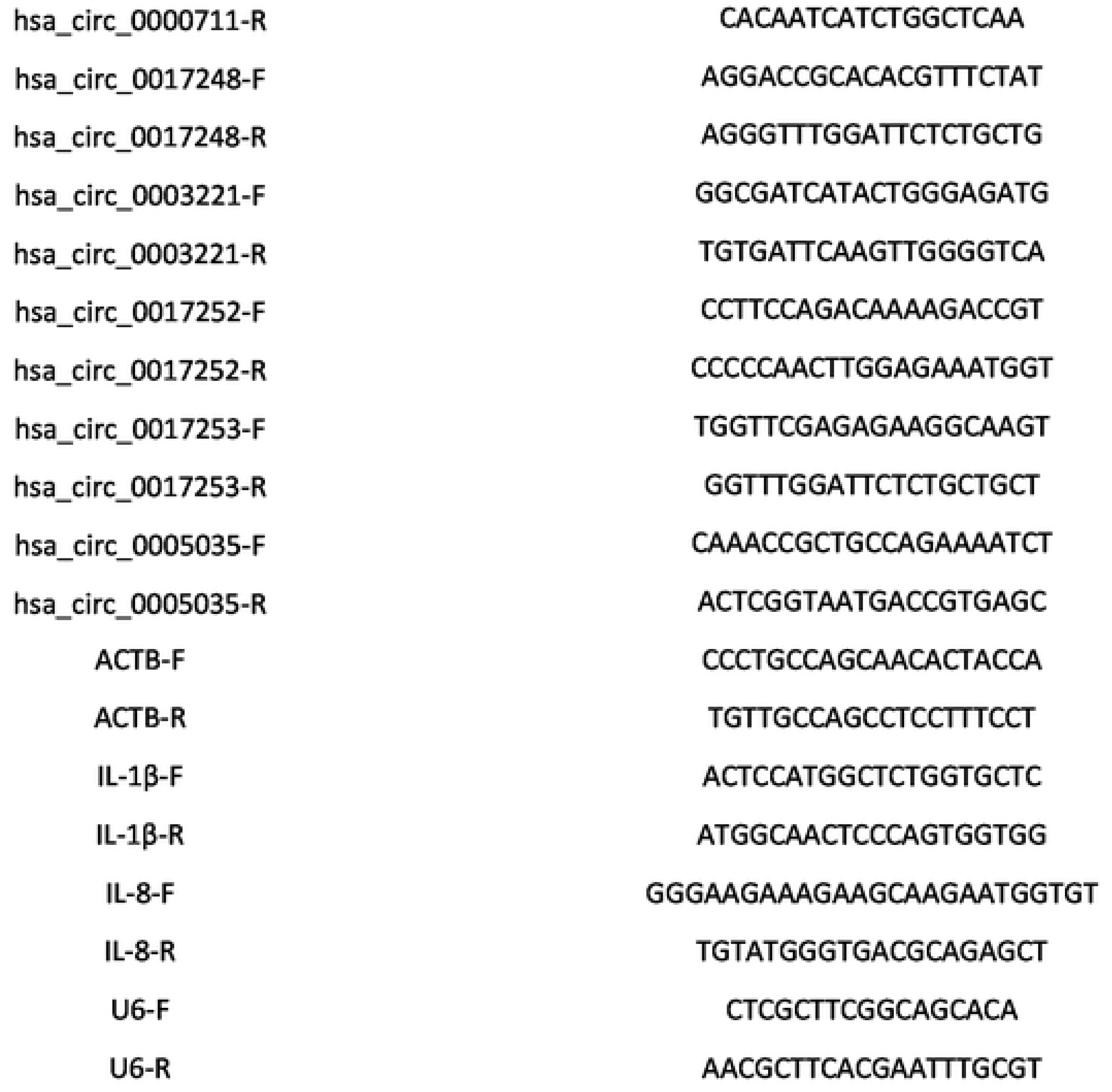
circRNA primer sequence list.

**TableS2.**
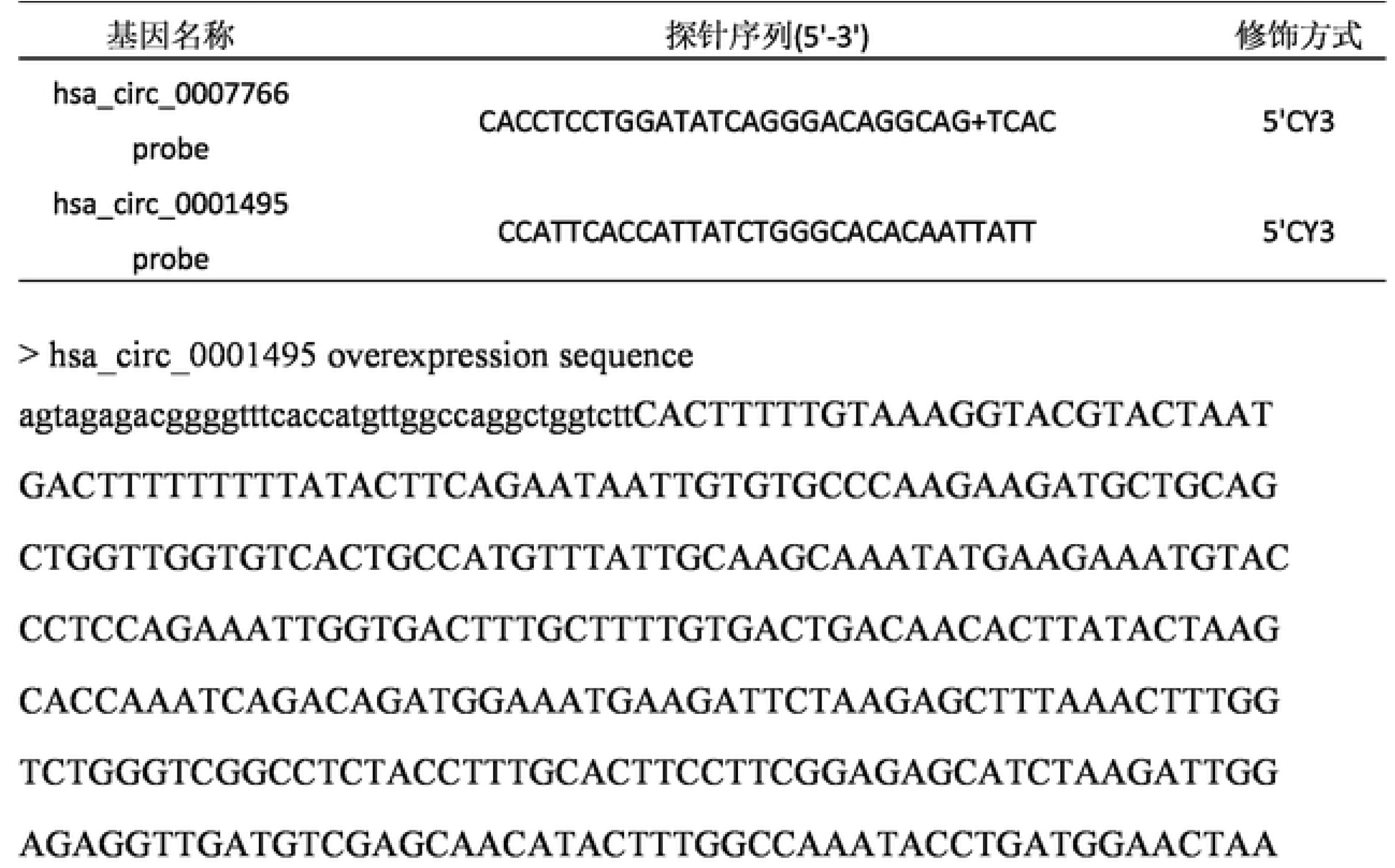
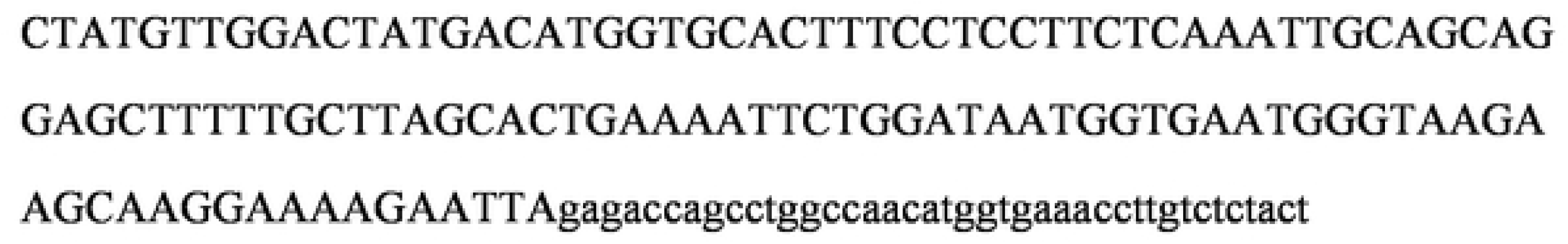
FISH probe sequence.

## Graphical Abstract

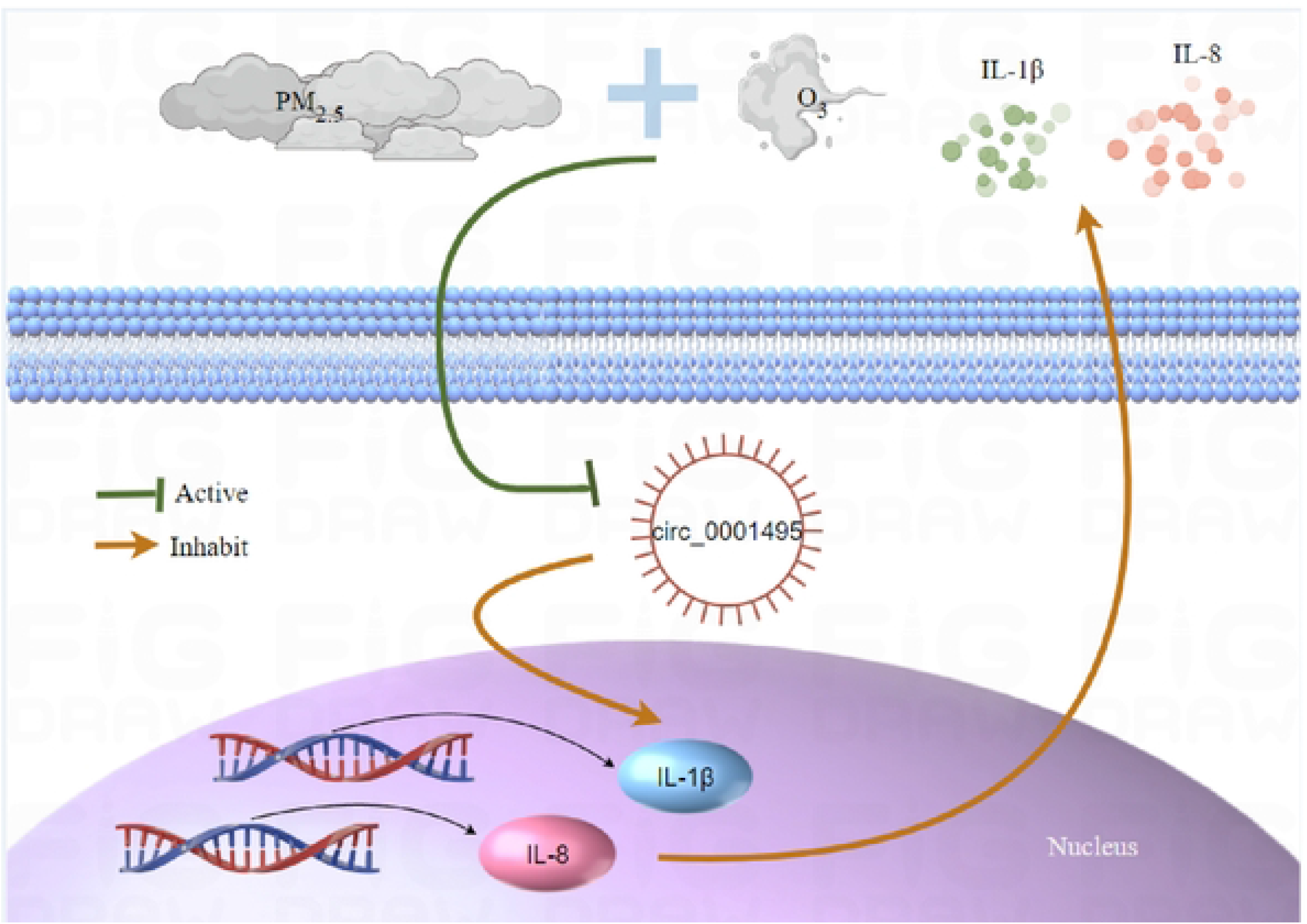

